# Testing a convolutional neural network-based hippocampal segmentation method in a stroke population

**DOI:** 10.1101/2020.01.28.924068

**Authors:** Artemis Zavaliangos-Petropulu, Meral A. Tubi, Elizabeth Haddad, Alyssa Zhu, Meredith N. Braskie, Neda Jahanshad, Paul M. Thompson, Sook-Lei Liew

## Abstract

As stroke mortality rates decrease, there has been a surge of effort to study post-stroke dementia (PSD) to improve long-term quality of life for stroke survivors. Hippocampal volume may be an important neuroimaging biomarker in post-stroke dementia, as it has been associated with many other forms of dementia. However, studying hippocampal volume using MRI requires hippocampal segmentation. Advances in automated segmentation methods have allowed for studying the hippocampus on a large scale, which is important for robust results in the heterogeneous stroke population. However, most of these automated methods use a single atlas-based approach and may fail in the presence of severe structural abnormalities common in stroke. Hippodeep, a new convolutional neural network-based hippocampal segmentation method, does not rely solely on a single atlas-based approach and thus may be better suited for stroke populations. Here, we compared quality control and the accuracy of segmentations generated by Hippodeep and two well-accepted hippocampal segmentation methods on stroke MRIs (FreeSurfer 6.0 whole hippocampus and FreeSurfer 6.0 sum of hippocampal subfields). Quality control was performed using a stringent protocol for visual inspection of the segmentations, and accuracy was measured as volumetric and spatial comparisons to the manual segmentations. Hippodeep performed significantly better than both FreeSurfer methods in terms of quality control and spatial accuracy. Overall, this study suggests that both Hippodeep and FreeSurfer may be useful for hippocampal segmentation in stroke rehabilitation research, but Hippodeep may be more robust to stroke lesion anatomy.

## Introduction

According to the World Health Organization, approximately 10.3 million people experience a stroke each year worldwide (Feigin et al., 2017). Post-stroke dementia (PSD), defined as any dementia occurring after stroke (including cognitive impairment, Alzheimer’s disease (AD), and vascular dementia) presents in roughly 30% of stroke survivors (Mok et al., 2017). PSD is one of the leading causes of dependency in stroke survivors (Leys et al., 2005) and is of growing concern for patients, families, and health-care providers as stroke survival rates improve (Dichgans, 2019). Therefore, early neuroimaging biomarkers that may contribute to PSD remain important to investigate.

The hippocampus may be an important biomarker for PSD. The hippocampus, essential for memory function, is vulnerable to pathology and atrophy in multiple dementia subtypes (Braak & Braak, 1991; Braskie & Thompson, 2014, Halliday, 2017), including PSD (Gemmel et al., 2012, Gemmel et al., 2014). The hippocampus is usually not directly impacted by an ischemic stroke lesion (Szabo et al., 2009). However, emerging evidence suggests that diaschisis, where stroke lesions can cause indirect effects on distant brain structures, may contribute to hippocampal atrophy (Klingbeil et al., 2020). Specifically, ischemic stroke is associated with reduced hippocampal volume, which is detectable *in vivo* by non-contrast MRI (Werden et al., 2017).

Robustly studying patterns of hippocampal atrophy after stroke requires large datasets, given the vast heterogeneity of stroke lesions in terms of lesion size, location, and presentation. This has incentivized large multi-center worldwide consortia to obtain large samples of post-stroke MRI to evaluate robust PSD hippocampal patterns. Consortia around the world - such as the Cognition and Neocortical Volume After Stroke Consortium (CANVAS; Brodtmann et al., 2014) and the Stroke and Cognition Consortium (STROKOG; Sachdev et al., 2017) - have made significant efforts to study the role of hippocampal volumes in the context of overall stroke recovery on a large scale. The Enhancing Neuroimaging through Meta-Analysis (ENIGMA) Stroke Recovery working group (Liew et al., 2020) is also interested in studying the post-stroke hippocampus in the context of sensorimotor recovery. Currently, manual segmentations are arguably the gold standard for analyzing hippocampal volume in MRI studies (Frisoni et al., 2015), but this approach is extremely time consuming and not feasible for large datasets such as these. Therefore, efforts to develop and test automated hippocampal segmentation methods have been undertaken to provide a more efficient way to study hippocampal volume on a large scale.

Current automated brain structure segmentation algorithms predominantly rely on atlas-based approaches, involving machine learning and sophisticated image registration to a single probabilistic atlas of pre-labeled regions. FreeSurfer (Fischl et al., 2002; Fischl, 2012), a robust method to segment both cortical and subcortical structures, is an atlas-based approach and commonly used to study hippocampal volume in cognitively healthy populations (Ritchie et al., 2018; Nobis et al., 2019) as well as in people with neurodevelopmental, psychiatric, and neurodegenerative conditions (Schmaal et al., 2016; Hibar et al., 2017; van Erp et al., 2017; Müller-Ehrenberg et al., 2018; Zhao et al., 2019). Recent studies by Khlif et al., (2019a, 2019b) compared automated hippocampal segmentation methods, such as the gross hippocampal segmentation available in FreeSurfer version 5.3, version 6.0, and the ‘sum of subfields’ segmentation available in FreeSurfer version 6.0, in stroke populations. Khlif et al., (2019a, 2019b) reported that the FreeSurfer version 6.0 ‘sum of subfields’ segmentation was among the most accurate methods for estimating hippocampal volume in healthy and ischemic stroke populations with lesions outside the hippocampus.

FreeSurfer was specifically designed to account for structural brain abnormalities common to AD and aging (Fischl, 2012), which share some overlapping features with stroke populations (Mok et al., 2017; Yousufuddin & Young, 2019); perhaps as a result, FreeSurfer has performed relatively well in stroke studies. However, large brain lesions are distinct to stroke patients and can introduce large alterations to the expected spatial distribution of brain structures, presenting a significant challenge to FreeSurfer. FreeSurfer, and most other probabilistic atlas-based automated segmentation methods, were not explicitly designed to accommodate significant brain injury pathology (Irimia et al., 2012) and are more likely to fail in the presence of large lesions (Yang et al., 2016). New methods that do not use single atlas-based automated segmentation methods may better accommodate stroke pathology and help improve segmentation accuracies in studies of the hippocampus in stroke. Related to this, recently, *Hippodeep*, a new convolutional neural network-based (CNN) algorithm, emerged as a fast and robust hippocampal segmentation method (Thyreau et al., 2018). Hippodeep relies on hippocampal ‘appearance’ instead of a single atlas-based approach. Hippodeep has better spatial agreement with manual segmentations than FreeSurfer version 6.0 ‘sum of subfields’ segmentation in healthy aging populations but has not yet been evaluated in a stroke population (Nogovitsyn et al., 2019).

Our study sought to expand on previous findings (Khlif et al. 2019a; Khlif et al., 2019b; Nogovitsyn et al., 2019) and evaluate how Hippodeep compares to previously tested methods for hippocampal segmentation in a stroke population. Using the Anatomical Tracings of Lesions After Stroke dataset (ATLAS; Liew et al., 2018), we compared Hippodeep, FreeSurfer version 6.0 gross hippocampal segmentation, and FreeSurfer version 6.0 ‘sum of subfields’ segmentation in terms of 1) quality control (QC) and 2) accuracy when compared to expert manual segmentations. QC and accuracy provide different but complementary evaluations of hippocampal segmentation. QC was done by visually inspecting segmentations to determine which segmentations failed to satisfy our predetermined criteria for a good quality segmentation. We measure accuracy by calculating two complementary measurements including 1) intra-class correlation, which is a measure of how similar the *volumes* are to their corresponding manual segmentations and 2) spatial overlap, which is a measure of the *spatial correspondence* between the labeled hippocampal voxels relative to the voxels of the corresponding manual segmentation. We hypothesized that Hippodeep’s CNN-based method would perform better on lesioned brain anatomy, resulting in fewer segmentation failures and more accurate hippocampal segmentations than either FreeSurfer method.

## Methods

### I. Data Acquisition

For our analyses, we used the ATLAS dataset (N=229), an open source dataset of anonymized T1-weighted structural brain MRI scans of stroke patients and corresponding manually traced lesion masks (Liew et al., 2018). All 229 scans were completed on 3-Tesla MRI scanners at a 1 mm isotropic resolution, intensity normalized and registered to the MNI-152 template space. T1-weighted MRIs, lesion masks, and metadata are publicly available for download (Liew et al., 2018). We analyzed the normalized data from these 229 participants as the input data to test the three automated segmentation methods.

### II. Hippocampal Segmentation Methods

#### II.a. FreeSurfer version 6.0

As mentioned previously, Khlif et al. (2019a, 2019b) found ‘sum of subfields’ segmentation available in FreeSurfer version 6.0 to be one of the best performing segmentation methods for the stroke data they evaluated. FreeSurfer is an atlas-based software that employs a Bayesian statistical approach to segment and label brain regions (Fischl, 2012). It involves a series of data preprocessing steps, such as intensity normalization, mapping of the input brain to a probabilistic brain atlas, estimation of statistical distributions for the intensities of different tissue classes, and labeling of cortical and subcortical structures based on known information on the locations and adjacencies of specific brain substructures (Fischl et al., 2002).

FreeSurfer version 6.0 can output segmentations of 13 hippocampal subregions using a refined probabilistic atlas (Fischl, 2012). This atlas was built from a combination of ultra-high resolution *ex vivo* and *in vivo* MRI scans, to identify borders between subregions of the hippocampus (Iglesias et al., 2015). The *ex vivo* scans included autopsy samples of participants with AD and controls scanned with a 7T scanner at 0.13 mm isotropic resolution that were then manually segmented by expert neuroanatomists. The *in vivo* data consisted of manual segmentations from 1mm isometric resolution T1-weighted MRI data acquired using a 1.5 T scanner from controls and participants with mild dementia. *In vivo* and *ex vivo* segmentations were combined to create one single computational atlas of hippocampal subfields. In this study, we combined the volumes of the individually labeled hippocampal subfields output by FreeSurfer version 6.0 to create a segmentation of the entire hippocampus, which we refer to as FS-Subfields-Sum throughout our study.

FreeSurfer also outputs a separate hippocampal segmentation using a different atlas, the Desikan-Killiany atlas (Desikan et al., 2006). The Desikan-Killiany atlas was built using 40 T1-weighted 1×1×1.5 mm spatial resolution MRIs acquired on a 1.5T scanner. These 40 participants were of ranging age and cognitive status with the intent to include a range of anatomical variance common to aging and dementia in the atlas. This hippocampal volume from the Desikan-Killiany atlas can be calculated using the hippocampus labels of the *aseg* FreeSurfer output file.

FreeSurfer outputs segmentations to a FreeSurfer specific image space. The FreeSurfer command, *mri_label2vol*, was used to transform the segmentation back to the original MNI space used in the input for both FreeSurfer versions segmentations. Segmentations from the *aseg* output are referred to as FS-Aseg throughout our study.

Prior studies have reported an inability to run FreeSurfer on certain participants with large lesions (Bigler et al., 2013; Khlif et al., 2019a). In an effort to generate the maximum number of segmentations, scans that were not segmented on the initial FreeSurfer analysis were run a second time through FreeSurfer.

#### II.b Hippodeep

Hippodeep is a recent automated hippocampal segmentation algorithm that has not yet been tested in stroke populations. Hippodeep does not warp individual images to an atlas; instead, it relies on a hippocampal appearance model learned from existing FreeSurfer v5.3 labeled online datasets as well as synthetic data (Thyreau et al., 2018). Two types of synthetic data are included in training the Hippodeep CNN. The first synthetic data is a manual segmentation of a synthetic high-resolution image of the hippocampus generated from an average of 35 variations of MRI scans of a single healthy participant. The purpose of segmenting the hippocampus on a high-resolution image (0.6 mm isotropic resolution) is to provide more detailed boundary information to the CNN that might not be as clear on a lower resolution image. The second type of synthetic data used to train the Hippodeep CNN are artificially geometrically distorted versions of the FreeSurfer v5.3 training data. Some of the distortion goes beyond the range of clinically plausible values but remains realistic enough to be easily delineated by a human rater. The purpose of this distorted data is to provide relevant training guidance to the CNN. By training the CNN on unconventional anatomy, Hippodeep may be more robust to severe stroke pathology. Details on the specifics of how the synthetic data were generated may be found in Thyreau et al. (2018).

Hippodeep outputs a probabilistic segmentation map calculated using a loss function to allow for the uncertainty of voxels along the perimeter of the hippocampus in native space, which can then be optionally thresholded. The probabilistic segmentation was converted to a binary mask of the hippocampus, as recommended by Thyreau et al., (2018).

#### II.c Manual Segmentations

We tested the accuracy of the automated methods by randomly selecting 30 participants for whom all three automated segmentation algorithms (FS-Aseg, FS-Subfields-Sum, and Hippodeep) were able to successfully output hippocampal segmentations. Only participants with unilateral lesions were considered. The ATLAS data was organized by lesion size and divided into thirds; small (range = 0.18 - 4.82 cubic centimeters (cc)), medium (range = 4.98 - 22.7cc), and large (range = 23.6 - 291.0cc) lesions. From each lesion size group, five participants with right hemisphere lesions and five participants with left hemisphere lesions were randomly selected. In this way, we examined the influence of lesion size across a broad range of lesion sizes, and with lesions equally distributed across hemispheres. In this data sample, all lesions occurred outside the medial temporal lobe.

Hippocampi for the subset of these 30 participants were manually traced by an expert rater (AZP), strictly adhering to the EADC-ADNI harmonized protocol for manual hippocampal segmentation (Boccardi et al., 2015; Frisoni et al., 2015). Coronal slices were used to trace the hippocampi using ITK-Snap (Yushkevich et al., 2006). The sagittal view was used to confirm hippocampal boundaries and edit the segmentations. Hippocampi were segmented blindly based on participant ID alone, starting with the left hippocampus, followed by the right hippocampus. Bilateral hippocampi were never overlaid on the T1-weighted image at the same time to avoid using the segmentation from one hemisphere to bias the other. The manual segmentations were checked for quality by another expert in hippocampal neuroanatomy (MAT). All manual segmentations are available for download here: https://github.com/npnl/Hippocampal_Segmentation

### III. Analyses

#### III.a Quality Control (QC)

We manually assessed the quality of segmentations produced by each automated hippocampal segmentation method in the full ATLAS dataset (N=229) using the ENIGMA Stroke Recovery QC protocol (Liew et al., 2020). Briefly, a trained researcher (AZP) reviewed nine slices of each brain (3 coronal, 3 axial, and 3 sagittal) with the bilateral segmentations overlaid on the T1, which were generated for each participant (**Supplementary Figure 1**). A segmentation failed QC if the segmentation grossly underestimated the hippocampus (underestimated), overestimated by including regions of the brain outside the hippocampus (overestimated), missed the hippocampus entirely (miss), or failed to output a segmentation (no output) (**Figure 1**).

**Figure 1.**
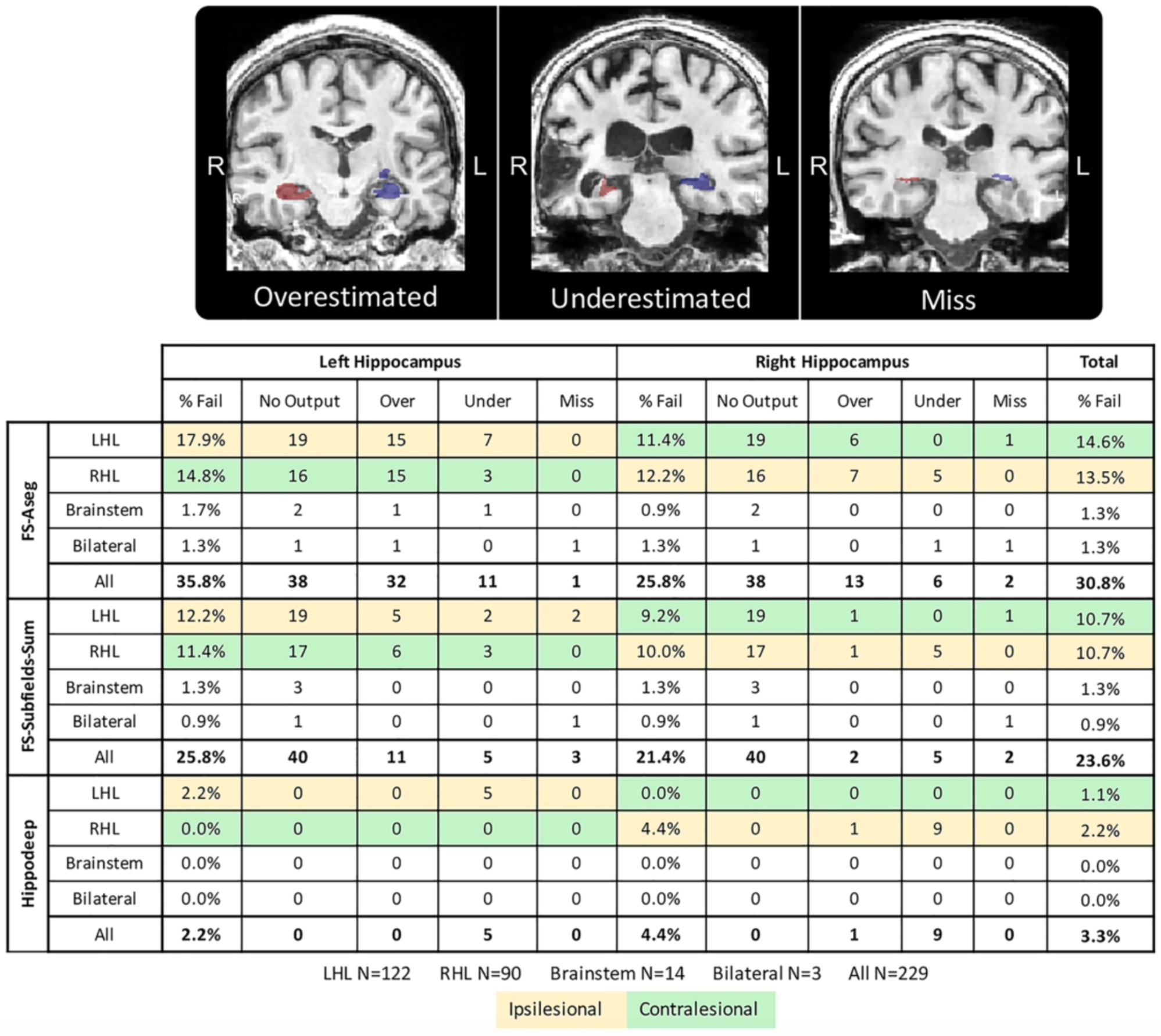
Hippocampal segmentations produced by automated segmentation methods (Hippodeep, FS-Aseg, and FS-Subfields-Sum) on the 229 ATLAS participants were inspected for quality according to the ENIGMA Stroke Recovery Working Group quality control (QC) protocol (Liew et al., 2020). Segmentations failed QC for four possible reasons: 1) failing to output a segmentation entirely (no output) 2) including voxels in the segmentation that are clearly outside of the hippocampus (overestimating) 3) underestimating the hippocampus (underestimating), or 4) producing a segmentation that misses the hippocampus entirely (miss). In this figure, we report the total breakdown of the QC results by hemisphere. The results are further broken down by location of lesion (LHL= left hemisphere lesion, RHL= right hemisphere lesion). Percent fail for left and right hippocampi is calculated as the total number of segmentations that failed QC for the specified hemisphere divided by 229. Percent fail for total is the number of segmentations divided by 458.

QC was reported in two levels of stringency: 1) methods-wise QC and 2) across-methods QC, similar to Sankar et al. (2017). For methods-wise QC, we calculated a QC fail rate for each automated segmentation method by dividing the total number of segmentations that failed for each segmentation method by the total number of segmentations (229 participants * 2 hippocampi = 458 segmentations).

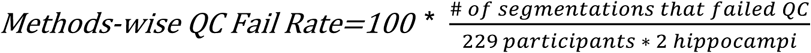

Across-methods QC fail rate was calculated as the total number of participants for which all three automated algorithms failed QC on at least one of the hippocampi divided by the total number of participants in the analysis (N=229).

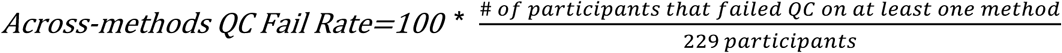

QC images and scores for each of the automated hippocampal segmentation methods on the 229 participants in ATLAS are available here: https://github.com/npnl/Hippocampal_Segmentation.

#### III.b Statistical Analysis of Accuracy

All statistical analyses were conducted in R-Studio version 1.1.463. To promote open science and reproducibility, all statistical analyses and code used for this study can be found here: https://github.com/npnl/Hippocampal_Segmentation

##### III.b.i Volume Correlation Analysis Comparison of Volumes

We evaluated the agreement in hippocampal volume across segmentation methods in the dataset of 30 participants, by calculating the Pearson’s correlation coefficient (R; Pearson, 1895) and the intra-class correlation coefficient (Shrout and Fleiss, 1979) in the ipsilesional and contralesional hippocampi separately. We predetermined the number of segmentation methods and we assumed no generalization to a larger population. Therefore, we assumed fixed judges for the intra-class correlation statistical analyses (ICC3).

##### III.b.ii Spatial Overlap

To estimate spatial segmentation accuracy, we measured the spatial overlap between the 30 manually traced hippocampi and each automated segmentation method (FS-Aseg, FS-Subfields-Sum, Hippodeep) using the Dice Coefficient (DC). DC was calculated for ipsilesional and contralesional hippocampi separately. DC is commonly used to validate segmentation algorithms in neuroimaging (Dice et al., 1945; Zou et al., 2004) and is defined as:

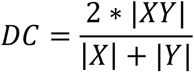

Here *X* represents the set of voxels in the manual segmentation and *Y* represents the set of voxels in the automated segmentation. DC can range from 0 (no spatial overlap) to 1 (complete overlap). For this study, we calculated DC using the flag *DiceandMinDistSum* of the ImageMath package from the Advanced Normalization Tools (ANTs) software (Avants et al., 2011).

A 2×3×3 Analysis of Variance (ANOVA) was performed to model dependencies of DC on the hemisphere with factors of lesion hemisphere (contralesional and ipsilesional), lesion size (small, medium, and large), and automated segmentation method (Hippodeep, FS-Subfields-Sum, FS-Aseg). A post-hoc paired *t*-test was used to compare DC between automated segmentation methods.

## Results

### I. Quality Control

First, we performed a rigorous quality control analysis for the segmentations generated by each automated method. This provided a sense of how robust each method was for generating good quality segmentations on the stroke data. The method-wise QC fail rate for FS-Aseg was 30.9% (N=144), 23.6% (N=108) for FS-Subfields-Sum, and 3.3% (N=15) for Hippodeep. The across-methods QC fail rate was 45.0% (N=103). A summary of reasons for QC fails by hemisphere for each segmentation algorithm can be found in **Figure 1**.

FS-Aseg did not output segmentations for 38 participants and FS-Subfields-Sum did not output segmentations for 40 participants (the 38 that did not output from FS-Aseg plus two additional participants). Of the 80 total hippocampi (40 participants * 2 hippocampi) that were not segmented by either FS-Aseg or FS-Subfields-Sum, 75 of these hippocampi were successfully segmented by Hippodeep and passed QC. Hippodeep also successfully segmented the remaining 5 hippocampi, but these did not pass QC and were all underestimated ipsilesional hippocampi. QC images of Hippodeep segmentations for participants who had no output by FS-Aseg or FS-Subfield-Sum are compiled in a file here: https://github.com/npnl/Hippocampal_Segmentation/Hippodeep_QC_for_no_output_FS_scans.pdf

### II. Accuracy

#### II.a Volume Correlation Analysis

In the subset of 30 participants with manually segmented hippocampi, we also compared hippocampal volume between automated and manual segmentations. All three segmentation methods overestimated both ipsilesional and contralesional hippocampal volume, compared to the manual gold standard (**Figure 2, Figure 3**). Hippodeep and FS-Subfields-Sum segmentations were not significantly different in volume (**Figure 3a**).

**Figure 2.**
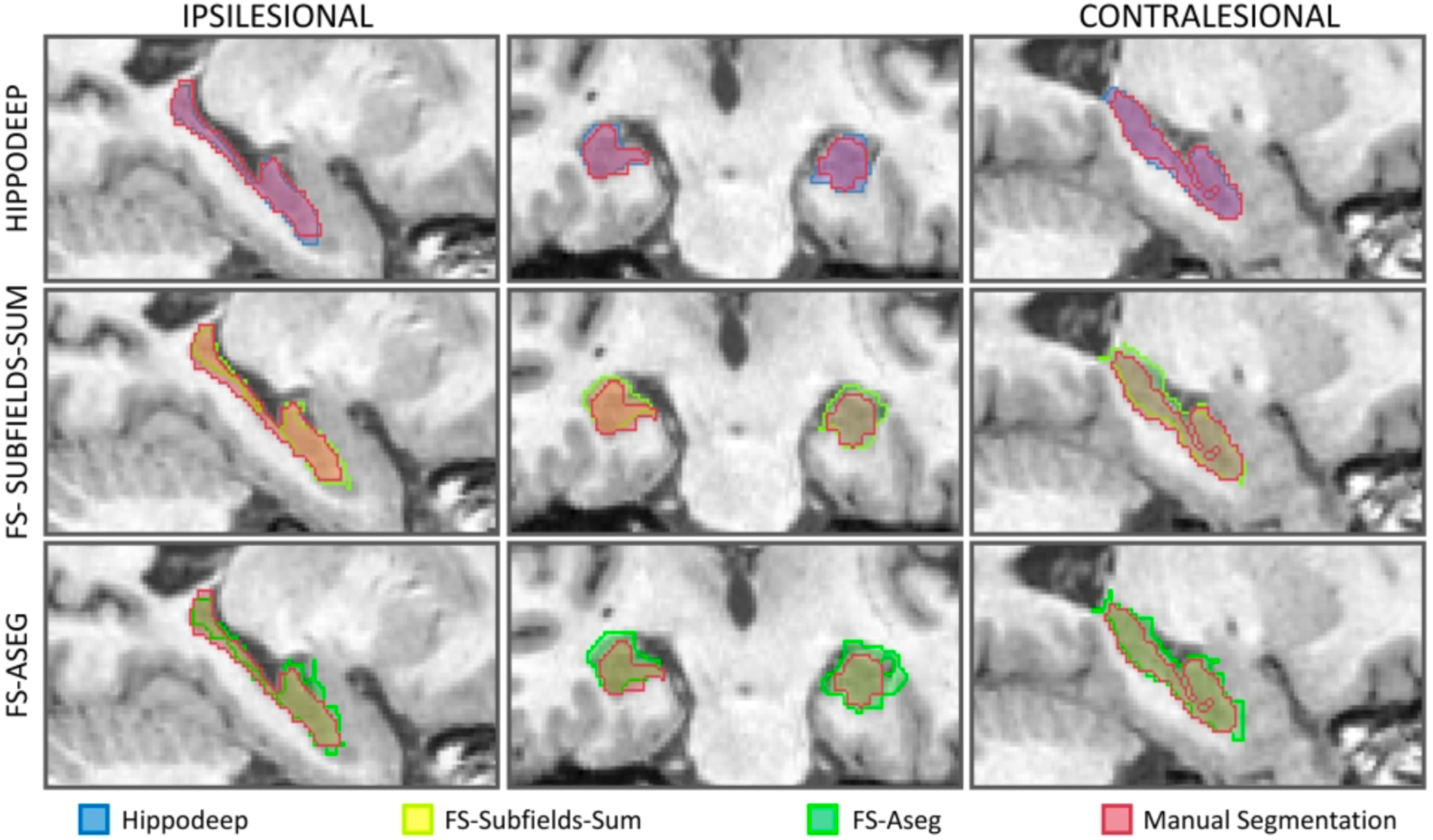
Automated hippocampal segmentations are overlaid, along with the manual segmentation, on MRI data from an example participant. Each row shows the results of a different automated segmentation method. The left column shows a sagittal view of the ipsilesional hemisphere, the middle column shows a coronal view of the body of bilateral hippocampi, and the rightmost column shows a sagittal view of the contralesional hemisphere.

**Figure 3.**
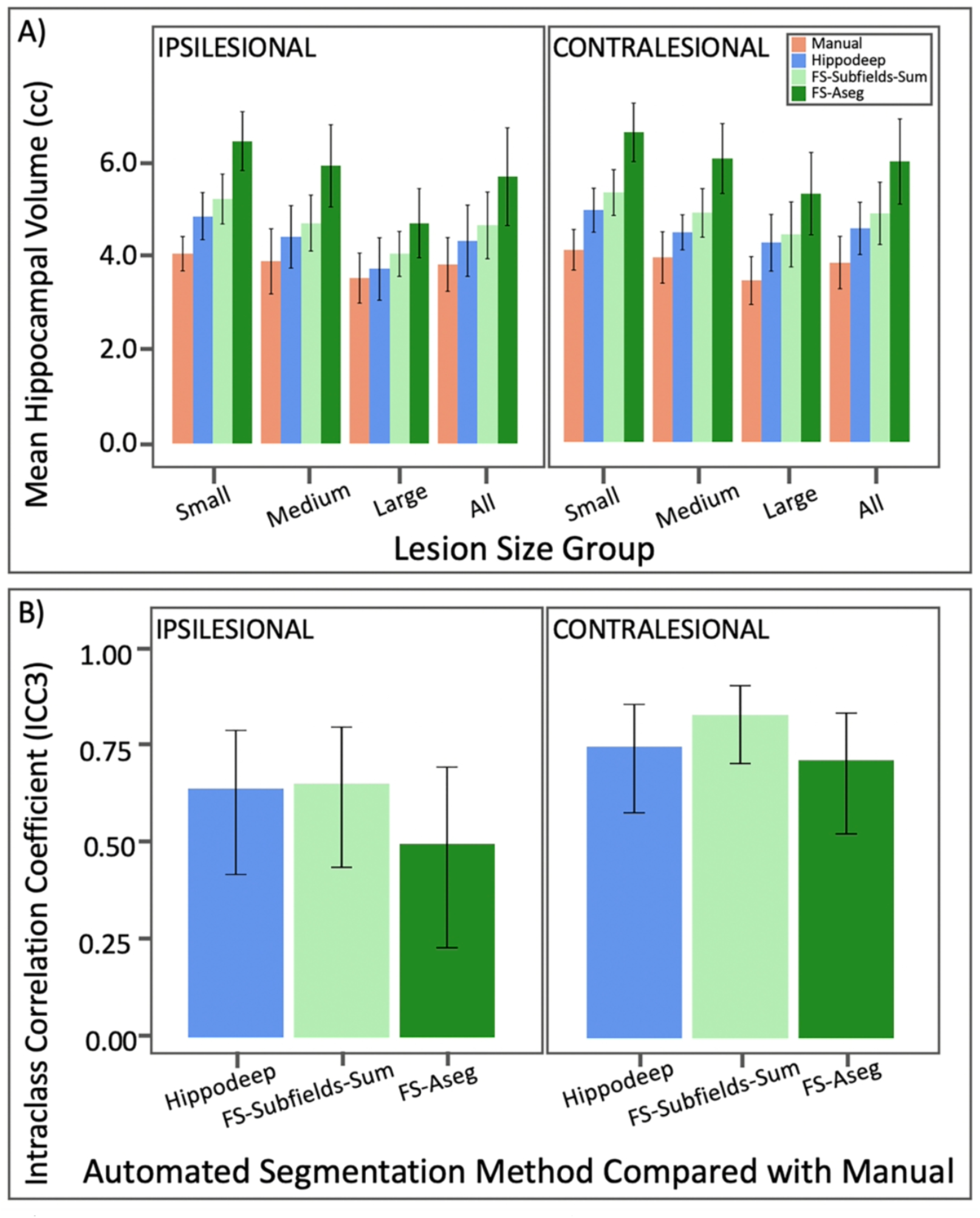
A) Mean hippocampal volume is plotted for manual and automated segmentation methods in the 30 participants with manually segmented hippocampi. All three automated segmentation methods on average overestimated the manually defined segmentation volume. This trend is consistently found for scans with small, medium, and large lesions. Error bars represent standard deviation. B) Intraclass Correlation Coefficient (ICC3) was calculated correlating volumes from each automated segmentation algorithm with manual segmentations. The error bars indicate the upper and lower bound of ICC3. FS-Subfields-Sum has the highest ICC3 with manual segmentations, although none of the ICC3 results are significantly different across automated methods.

As expected, volumes from all three segmentation methods were strongly correlated with volumes from the manual segmentations (**Table 1**). Volumes from FS-Subfields-Sum had the strongest correlation with manual segmentation volumes (ipsilesional ICC3 = 0.65; contralesional ICC3 = 0.83). Hippodeep measures were also strongly correlated with manual segmentation volumes (ipsilesional ICC3 = 0.64; contralesional ICC3 = 0.75). FS-Aseg was the least correlated with the manual segmentation volumes (ipsilesional ICC3 = 0.50; contralesional ICC3 = 0.71). Volumes from FS-Subfields-Sum and Hippodeep were strongly correlated with each other (ipsilesional ICC3 = 0.91; contralesional ICC3 = 0.90). However, upper and lower bounds for ICC3 indicated there were no significant differences among ICC3 values across comparisons (Figure 3b). Volumes from all three segmentation methods were strongly correlated with each other (**Table 2**).

**Table 1.** Intraclass correlation coefficient (ICC3), Pearson’s correlation coefficient (R), and *p*-values were calculated correlating hippocampal volume from the automated segmentation methods to the manual segmentations. Correlations between FS-Subfields-Sum and Hippodeep are also shown because the resulting hippocampal volumes were very similar.

|  | Ipsilesional |  |  | Contralesional |  |  |
| --- | --- | --- | --- | --- | --- | --- |
|  | ICC3 | R | *p*-value | ICC3 | R | *p*-value |
| FS-Aseg vs. Manual | 0.50 | 0.59 | 6.45 x10<sup>-4</sup> | 0.71 | 0.80 | 1.21 x10<sup>-7</sup> |
| FS-Subfields-Sum vs. Manual | 0.65 | 0.67 | 5.71 x10<sup>-5</sup> | 0.83 | 0.84 | 5.38 x10<sup>-9</sup> |
| Hippodeep vs. Manual | 0.64 | 0.69 | 2.18 x10<sup>-5</sup> | 0.75 | 0.75 | 1.91 x10<sup>-6</sup> |

**Table 2.** Intraclass correlation coefficient (ICC3) was calculated to compare segmentations from each automated method. Upper and lower boundaries of ICC3 are also reported. The volumes for all hippocampal segmentations are highly correlated, implying inter-algorithm consistency.

| Method | Ipsilesional |  |  | Contralesional |  |  |
| --- | --- | --- | --- | --- | --- | --- |
|  | ICC3 | Lower Bound | Upper Bound | ICC3 | Lower Bound | Upper Bound |
| FS-Subfields-Sum Vs. Hippodeep | 0.91 | 0.83 | 0.95 | 0.90 | 0.82 | 0.94 |
| FS-Subfields-Sum vs FS-Aseg | 0.89 | 0.80 | 0.94 | 0.92 | 0.85 | 0.96 |
| FS-Aseg vs Hippodeep | 0.85 | 0.74 | 0.92 | 0.79 | 0.64 | 0.88 |

#### II.b Spatial Overlap (DC Analysis)

We calculated the Dice Coefficient for each automated segmentation compared to the manual segmentation to assess the accuracy of each method in the dataset of 30 participants. This provides a quantitative assessment of the spatial overlap between the automated and manual segmentations. We performed an ANOVA to evaluate how DC differed across automated segmentation method, lesioned hemisphere, and lesion size. The ANOVA revealed that segmentation method was the only significant factor for differences in DC (F = 402.3; *p*-value < 2 ×10^−16^). Hippodeep had the highest average DC (ipsilesional = 0.84 ± 0.03; contralesional = 0.84 ± 0.02), followed by FS-Subfields-Sum (ipsilesional = 0.73 ± 0.03; contralesional = 0.72 ± 0.03), followed by FS-Aseg (ipsilesional = 0.69 ± 0.04; contralesional = 0.68 ± 0.03). We performed a paired t-test to see if DC significantly differed between the automated segmentation methods. Results from the *t*-test showed that the DC for Hippodeep was significantly higher than the DC for FS-Subfields-Sum (ipsilesional *p*-value *=* 1.03 ×10^−17^, *t*-value*=* 18.7; contralesional *p*-value *=* 4.68 ×10^−21^, *t*-value *=* 24.8) and FS-Aseg (ipsilesional *p*-value *=* 3.34 ×10^−21^, *t*-value *=* 25.1; contralesional *p*-value *=* 1.05 ×10^−23^, *t*-value *=* 30.8) (**Figure 4; Table 3**).

**Table 3.** Here, we show the results from the post-hoc t-tests comparing Dice Coefficients between automated methods.

|  | Ipsilesional |  | Contralesional |  |
| --- | --- | --- | --- | --- |
|  | *p*-value | *t*-value | *p*-value | *t*-value |
| Hippodeep vs. FS-Subfields-Sum | $1.03 \times 10^{-17}$ | 18.7 | $4.68 \times 10^{-21}$ | 24.8 |
| Hippodeep vs. FS-Aseg | $3.34 \times 10^{-21}$ | 25.1 | $1.05 \times 10^{-23}$ | 30.8 |
| Subfields-Sum vs. FS-Aseg | $3.68 \times 10^{-12}$ | 11.3 | $2.17 \times 10^{-15}$ | 15.3 |

**Figure 4.**
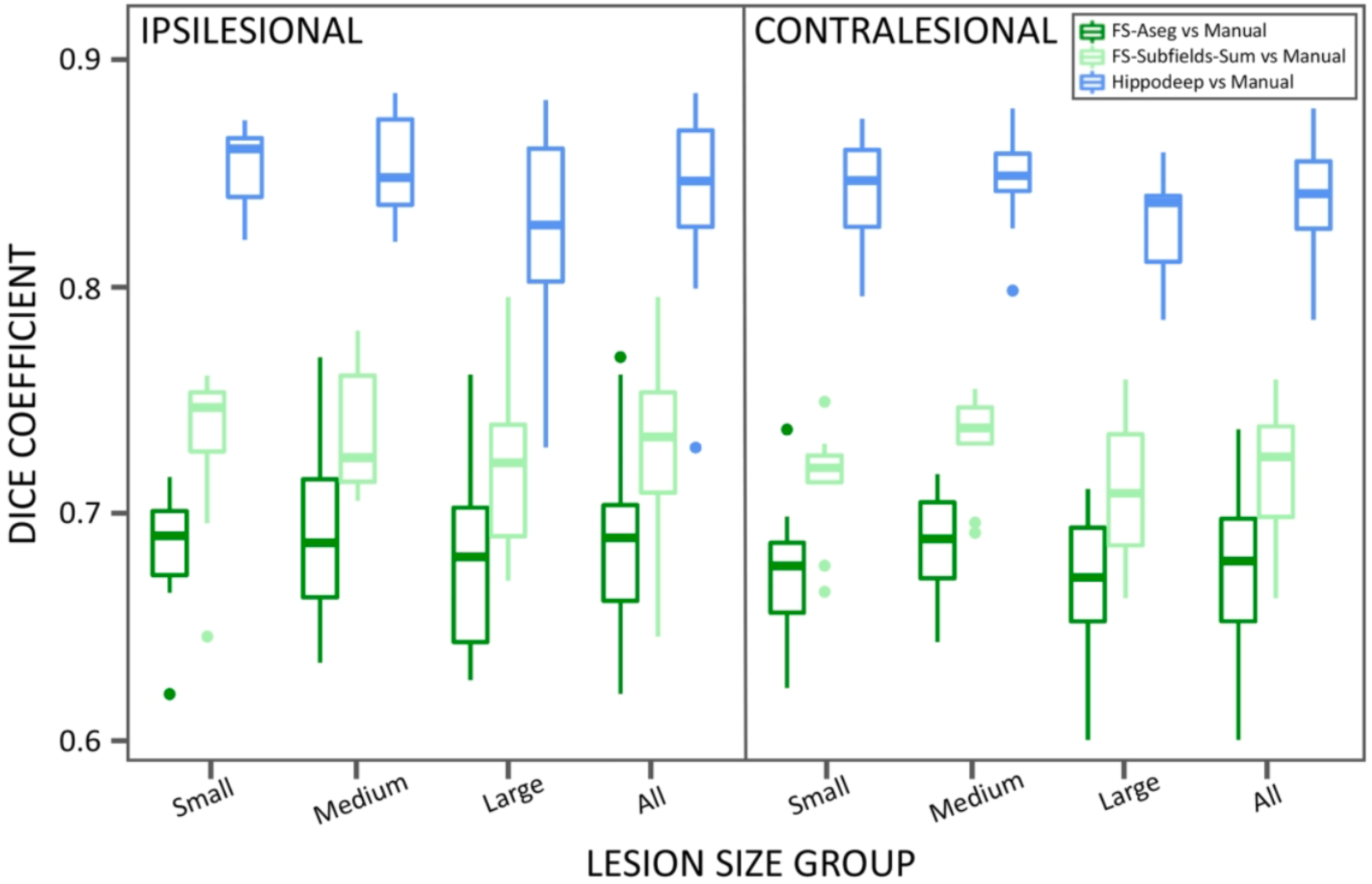
Average Dice Coefficient (DC) for Hippodeep vs. Manual segmentations was significantly higher than FS-Subfields-Sum vs. Manual (ipsilesional *p*-value *=* 1.03 ×10^−17^, *t*-value*=* 18.7; contralesional *p*-value *=* 4.68 ×10^−21^, *t*-value *=* 24.8) and FS-Aseg vs. Manual (ipsilesional *p*-value *=* 3.34 ×10^−21^, *t*-value *=* 25.1; contralesional *p*-value *=* 1.05 ×10^−23^, *t*-value *=* 30.8). DC did not differ significantly across lesion size.

## Discussion

In this study, we compared the quality control (QC) and accuracy of three automated segmentation algorithms (Hippodeep, FS-Subfields-Sum, FS-Aseg) used to estimate hippocampal volume in individuals with stroke. Our study found that Hippodeep was able to generate the greatest number of segmentations that passed QC and performed the best in terms of accuracy, as measured by the Dice Coefficient, while FS-Subfields-Sum performed slightly higher in terms of intraclass correlations (ICC3). This suggests that, while both Hippodeep and FS-Subfields-Sum produce good correspondence with manual hippocampal segmentations, Hippodeep is more accurate in terms of the actual voxels identified as the hippocampus while FS-Subfields-Sum produces a closer match on the overall volume of the hippocampus.

Hippodeep had the smallest methods-wise QC fail rate of the three automated segmentations tested (3.3%). FS-Subfields-Sum had the second lowest methods-wise QC fail rate (23.6%) followed by FS-Aseg (30.8%). Sankar et al., (2017) report high rates of poor-quality hippocampal segmentation across multiple automated segmentation algorithms, including FS-Aseg in elderly populations. While a certain amount of segmentation failure is expected for automated methods, automated segmentations in stroke populations are challenged by stroke pathology. An estimated 10-20% of FreeSurfer subcortical segmentations do not pass quality control in the ENIGMA Stroke Recovery Working Group data (Liew et al., 2020). In the stroke data we used here, Hippodeep was able to generate segmentations of adequate quality for 27.5% more hippocampi than FS-Aseg and 20.3% more than in FS-Subfields-Sum. Hippodeep generated volume estimates for all of the participants whose data could not be run through FreeSurfer in our study, and all but 5 of these segmentations passed QC. Therefore, Hippodeep can potentially help to maximize the number of participants included in analyses whose data might not run successful through FreeSurfer, potentially boosting statistical power, and reducing the bias that can come from excluding participants. Obtaining robust statistical power is of keen interest to the stroke recovery field, as a recent review by Kim & Winstein (2017) found that less than 30% of stroke recovery studies met the appropriate sample size criteria to achieve sufficient statistical power for predicting recovery. Our understanding of the role of hippocampal volume in stroke recovery will benefit from studies with larger, more representative samples.

In addition, hippocampal segmentation using Hippodeep resulted in the highest similarity index (DC) to manual tracing, indicating a high level of segmentation accuracy. Hippodeep performed significantly better in terms of DC than FS-Subfields-Sum and FS-Aseg. FS-Subfields-Sum also yielded a high similarity index to manual segmentations, consistent with prior results (Khlif et al. 2019a; Khlif et al., 2019b). Mean DC values for both Hippodeep and FS-Subfields-Sum were above a DC of 0.7, which is a recommended threshold for DC to indicate good segmentation overlap (Zou et al., 2004).

DC was influenced by segmentation method, but not by lesion size or lesion hemisphere. There was also a wide range in lesion sizes for scans that failed to output FS-Aseg and FS-Subfields-Sum segmentations. Although lesion size may not have a significant effect on DC in this sample of 30 participants, it may influence the segmentation accuracy and merits further investigation.

FS-Subfields-Sum and Hippodeep were both very competitive in terms of their correlations with ipsilesional and contralesional volume estimates. FS-Subfields-Sum segmentations were more highly correlated with manual segmentations (ICC3) than Hippodeep, although both were high and considered very reliable (Koo & Li, 2016). There were no significant differences across methods in ICC3 results. Ipsilesional FS-Aseg volume estimates had the lowest ICC3, but this correlation was still high enough to be considered moderately reliable (Koo & Li, 2016). For all three automated methods, contralesional ICC3 was higher than ipsilesional ICC3. Hippodeep and FS-Subfields-Sum may have performed better than FS-Aseg in terms of volumetric accuracy because information from a high-resolution hippocampus is included in both the Hippodeep and FS-Subfield-Sum algorithms. FS-Subfields-Sum is based on an atlas generated using manual segmentations on an ultra-high resolution atlas (0.13 mm isotropic resolution; Iglesias et al., 2015). Hippodeep uses information from a manually traced hippocampus on a synthetic high-resolution image (0.6mm isotropic resolution; Thyreau et al., 2018). In contrast, FS-Aseg uses the Desikan-Killiany atlas, which was built using only scans of 1 ×1 ×1.5mm resolution (Desikan et al., 2006). The Desikan-Killiany atlas was designed to segment many structures across the brain, many of which are clearly delineated on low resolution scans. Including more detailed information on hippocampal boundaries that appear ambiguous on a low-resolution MRI may improve segmentation performance. Further exploration of methodological aspects of successful automated segmentation methods may be helpful to inform future development of methods in populations with irregular neuroanatomy.

Beyond QCfailure rates and accuracy, there are other technical aspects to consider when comparing Hippodeep, FS-Subfields-Sum, and FS-Aseg. Hippodeep requires less computational power than FreeSurfer and runs within minutes, whereas FreeSurfer can take over 24 hours on a typical CPU (Thyreau et al., 2018; Nogovitsyn et al., 2019). However, Hippodeep only outputs estimates of the hippocampus and total intracranial volume. In addition to FS-Subfields-Sum and FS-Aseg, FreeSurfer also estimates other brain measures beyond hippocampal volume and intracranial volume, such as individual subfield volumes (Iglesias et al., 2015), and cortical and subcortical volumes, as well as thickness measures, and other vertex based measures and attributes that can be used for surface-based statistical analyses (Fischl, 2012). Additionally, FreeSurfer has an extensive archive of user questions for troubleshooting, while Hippodeep is a recent method that is not as extensively documented. Therefore, selection of the appropriate hippocampal segmentation method should be evaluated within the context of the study requirements and constraints (**Table 4**).

**Table 4.** Here we present a roadmap for using each of the automated hippocampal segmentation method tested in this paper. Study requirements and constraints should be considered when selecting which method to apply.

|  | How to Run | Mean Dice | Pros | Cons | Anatomical Variability |
| --- | --- | --- | --- | --- | --- |
| FS-Aseg | https://surfer.nmr.mgh.harvard.edu/<br>`recon-all -s subject`<br>Use aparc+aseg.mgz in subject/mri/ folder to extract label 17 for the left hippocampus and label 53 for the right hippocampus | Ipsi=<br>$0.69 \pm 0.04$<br>Contra=<br>$0.68 \pm 0.03$ | -Widely used in the literature<br>-Provides brain measures beyond the hippocampus (cortical and subcortical volumes and thickness, etc; Fischl, 2012)<br>-Extensive support archive (https://surfer.nmr.mgh.harvard.edu/fswiki/FreeSurferSupport) | -Did not output segmentations for all of the stroke data<br>-Modest accuracy with manual segmentation<br>-Time consuming<br>-Resource intensive<br>-Low resolution atlas | -Atlas created on 1x1x1.5 mm resolution scan<br>-Fimbria is included |
| FS-Subfields-Sum | https://surfer.nmr.mgh.harvard.edu/<br>`recon-all -s subject -hippocampal-subfields-T1`<br>rh.hippoSfLabels-T1.v10.FSvoxelSpace_native.mgz and lh.hippoSfLabels-T1.v10.FSvoxelSpace_native.mgz in subject/mri folder | Ipsi=<br>$0.73 \pm 0.03$<br>Contra=<br>$0.72 \pm 0.03$ | -Strong accuracy with manual segmentation<br>-Provides brain measures beyond the hippocampus (cortical and subcortical volumes and thickness, etc; Fischl, 2012)<br>-Provides information about individual subfield volumes (Iglesias et al. 2015)<br>-Widely used in the literature<br>-Extensive support archive (https://surfer.nmr.mgh.harvard.edu/fswiki/FreeSurferSupport) | -Did not output segmentations for all of the stroke data<br>-Time consuming<br>-Resource intensive | -Atlas created on 0.13 mm isotropic resolution scan<br>-Fimbria is included |
| Hippodeep | https://github.com/bthyreau/hippodeep<br>`deepseg1.sh subject_t1.nii.gz`<br>example_brain_t1_mask_L.nii.gz and example_brain_t1_mask_R.nii.gz | Ipsi=<br>$0.84 \pm 0.03$<br>Contra=<br>$0.84 \pm 0.02$ | -Strong accuracy with manual segmentation<br>-Can help maximize sample size<br>• Output segmentations for all of the stroke data<br>• Able to produce good segmentations for most of the participants that could not run through FreeSurfer<br>-Short run-time | -Only outputs hippocampal segmentations and total brain volume<br>-Not trained specifically on stroke data<br>-Newer method with limited support archives | -Includes a manual segmentation built on 0.6 mm isotropic resolution scan<br>-Fimbria is not included |

### Limitations

We note that a Dice Coefficient analysis only works as a validation metric to compare methods that aim to produce the same segmentation boundaries. A key methodological limitation to consider when comparing these segmentation methods is that each segmentation method has slightly different criteria for determining hippocampal boundaries (Desikan et al., 2006; Frisoni et al., 2015; Thyreau et al., 2018). Certain hippocampal boundaries (specifically along the head and the tail) are only visible in high-resolution MRIs and consensus on rules for delineating the boundaries of these regions on an MRI has not yet been reached (Olsen et al., 2019). Variability in image resolution used to develop each algorithm may also contribute mild variability in anatomical boundaries (**Table 4**). Additionally, Hippodeep uses the fimbria of the hippocampus as a boundary and does not include it in the segmentation while FS-Aseg and FS-Subfields-Sum both include this region. The HARP EADC protocol used to perform the manual segmentations does not include the fimbria. Therefore, some mild variability in the resulting correlations and spatial overlaps with accuracy versus manual segmentations is expected.

Another key methodological limitation to consider when comparing these segmentation methods is that none of these approaches were designed specifically to accommodate severe stroke pathology. The default atlases used in FreeSurfer, including FreeSurfer subfields, were created based on data from cognitively healthy elderly adults and patients with early AD pathology (Desikan et al., 2006; Iglesias et al., 2015). Stroke pathology, such as large lesions, hydrocephalus *ex vacuo* of the lateral ventricle (Nelson, 2003), and midline shifts (Liao et al., 2018), can alter expected spatial distribution of brain anatomy. As a result, stroke pathology can interfere with templates used by existing atlas-based approaches, resulting in inaccurate hippocampal segmentations. Although the CNN used in Hippodeep was not trained on data with stroke pathology, it is trained to anticipate extreme anatomical variability from the synthetic data. Being robust to extreme anatomical variability may explain why Hippodeep was able to perform well in stroke participants. Stroke-specific CNN hippocampal segmentation models that include stroke pathology in training data may further improve automated hippocampal segmentation in this population.

### Conclusion

In this study, we demonstrated that most robust hippocampal segmentation method (Hippodeep) also provided the most accurate segmentations. Hippodeep had the lowest method-wise QC fail, suggesting it may be the most robust to post-stroke anatomical distortions. The use of more accurate automated hippocampal segmentation methods may reveal clinical associations that are so far undetected. Additionally, future work should aim to extract subfields from the Hippodeep segmentation to further enhance our understanding of how the specific regions of the hippocampus are indirectly impacted by stroke lesions. Overall, our results suggest that Hippodeep may be an optimal method for accurate and robust hippocampal segmentation methods in diverse stroke populations.

## Data Availability

To promote open science and reproducibility, images used for quality control, manual segmentations, and all statistical analyses and code used for this study can be found here: https://github.com/npnl/Hippocampal_Segmentation. Any issues or feedback can be submitted on this page under “issues” on the Github system and a team of researchers will address these in a timely manner.

## Acknowledgments

This work was supported in part by an NIH K01 HD091283 (S.-L. Liew) and R01 NS115845 (S.-L. Liew), the NIH Big Data to Knowledge (BD2K) program U54 EB020403 (P. Thompson), P41 EB015922 (P. Thompson, N. Jahanshad), R01 AG059874 (N. Jahanshad), and F31 AG059356 (M. Tubi).

