## Supplementary Figure 1 for "Testing a convolutional neural network-based hippocampal segmentation method in a stroke population"

**Supplementary Material**

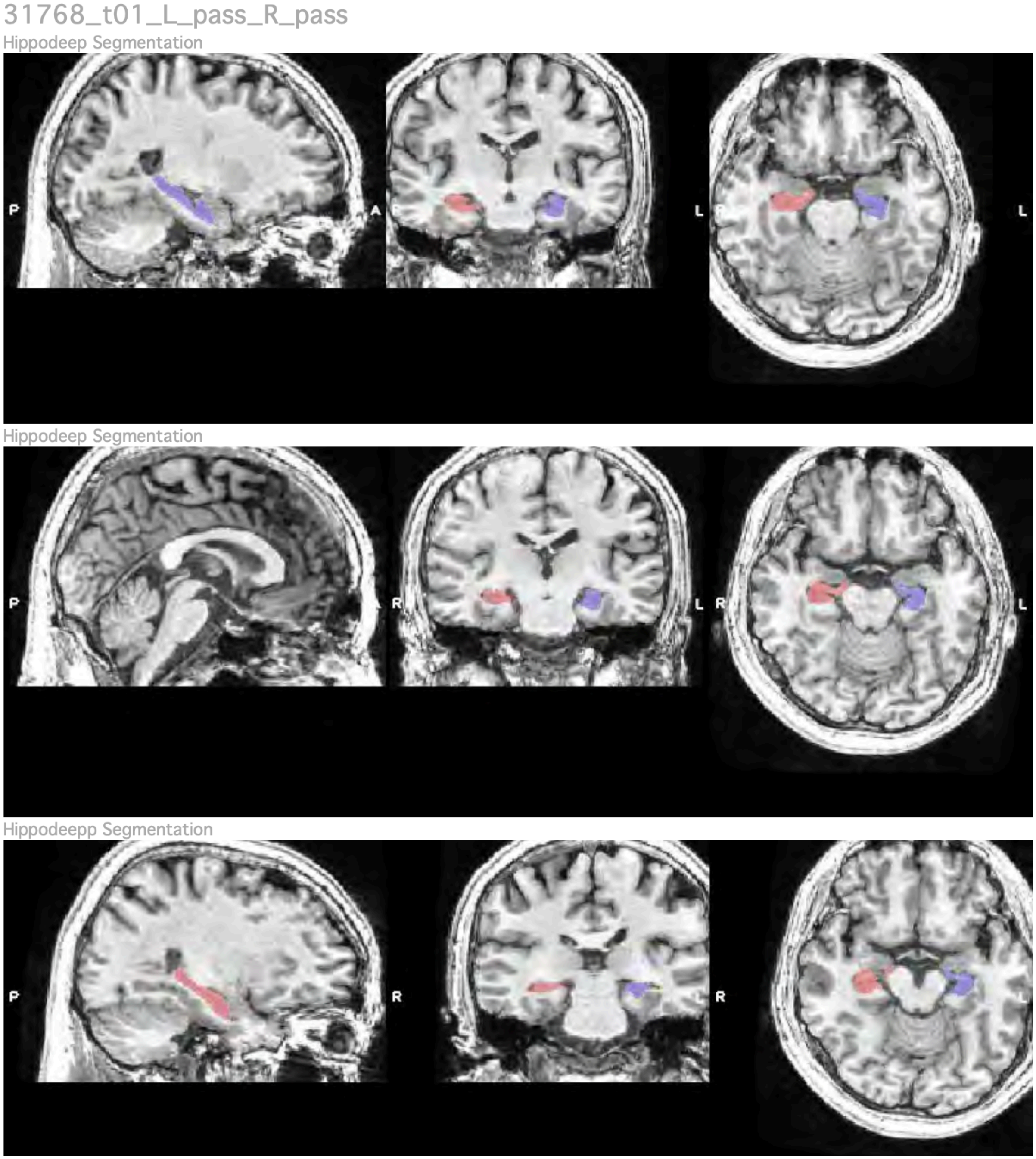


**Supplementary Figure 1:** Here we show an example of the 9 images per MRI used to perform Quality Control (QC). QC was done using 9 slices of the brain to observe the bilateral hippocampus. The right (*red*) and left (*blue*) hippocampi are overlaid onto the T1 and visually checked for quality. A segmentation failed QC if the segmentation grossly underestimated the hippocampus (underestimated), overestimated by including regions of the brain outside the hippocampus (overestimated), missed the hippocampus entirely (miss), or failed to output a segmentation (no output). The QC images for all 229 ATLAS scans can be found here: *https://github.com/npnl/Hippocampal_Segmentation.*
